# Dynamics of drug delivery determines course of evolution of antibiotic responses in bacteria

**DOI:** 10.1101/2023.11.29.569327

**Authors:** John C. Crow, Hao Geng, Timothy J. Sullivan, Shannon M. Soucy, Daniel Schultz

**Affiliations:** Department of Microbiology & Immunology, Dartmouth – Geisel School of Medicine, Hanover, NH 03755, USA; Department of Biomedical Data Science, Dartmouth – Geisel School of Medicine, Hanover, NH 03755, USA

## Abstract

To adjust to sudden shifts in conditions, microbes possess regulated genetic mechanisms that sense environmental challenges and induce the appropriate responses. The initial evolution of microbes in new environments is thought to be driven by regulatory mutations, but it is not clear how this evolution is affected by how quickly conditions change (i.e. dynamics). Here, we perform experimental evolution on continuous cultures of tetracycline resistant *E. coli* in different dynamical regimens of drug administration. We find that cultures evolved under gradually increasing drug concentrations acquire fine-tuning mutations adapting an alternative efflux pump to tetracycline. However, cultures that are instead periodically exposed to large drug doses evolve transposon insertions resulting in loss of regulation of the main mechanism of tetracycline resistance. A mathematical model shows that sudden drug exposures overwhelm regulated responses, which cannot induce resistance fast enough. These results help explain the frequent loss of regulation of resistance in clinical pathogens.

## Introduction

To enable the colonization of ecological niches where conditions change frequently^1,2^, microbes are equipped with inducible mechanisms that sense changes in their surroundings and initiate the appropriate cellular programs^3^. In some cases, responses are not time sensitive and mostly tune gene expression to optimal levels, such as when cells respond to shifts in nutrient conditions^4,5^. However, when cells are exposed to antibiotics or other harmful compounds, cell survival depends on the quick deployment of its defenses, while gene expression is still possible^6,7^. Therefore, even when microbes carry mechanisms to deal with hostile environments, sudden and frequent changes in conditions still threaten cells that are too slow to respond.

Dynamic environments, where conditions shift rapidly, pose fundamentally different selective pressures on the evolution of antibiotic responses^8–10^. Antibiotic resistance mechanisms originate in the soil, where antibiotic concentrations are typically low^11^, within a selection window that inhibits the growth of sensitive strains while enriching resistant subpopulations carrying beneficial mutations^12,13^. In contrast, antibiotics are used in the clinic in high doses, with the intent of wiping out entire microbial populations^14,15^. Such extreme environments pose strict bottlenecks, resulting in selective sweeps of any surviving mutants^16,17^, however unfit^18,19^. However, while much attention has been devoted to the evolution of antibiotic resistance under steady drug concentrations^20–22^, we still lack an understanding of how evolution proceeds in dynamic environments, where drug concentrations are high and change quickly. Recent studies suggest that the short-term evolution of microbes in such challenging environments relies heavily on mutations affecting regulatory pathways^23–25^.

To study the evolution of antibiotic responses, we focus on tetracycline resistance in *Escherichia coli*, which is mediated by two inducible efflux mechanisms capable of transporting the drug out of the cell - the *tet* and *acr* operons^26,27^ (Fig. 1A). While the *acr* operon transports a wide variety of toxic compounds and is part of the *E. coli* core genome^28,29^, the *tet* operon is a tetracycline specific module of the *E. coli* pan-genome that provides the bulk of resistance in strains where it is present^30,31^. Both mechanisms are regulated by repressors of the same family – TetR and AcrR, respectively^32,33^ – that can bind tetracycline and lose affinity for DNA, releasing expression of their respective efflux pumps, *tetA* and *acrAB*. Since the *acr* operon is not optimized for tetracycline efflux, several mutations in *acrB* have been reported to increase tetracycline resistance^34,35^. Additionally, active tetracycline transport via TetA involves an ion exchange that has been shown to disrupt the membrane potential, thereby posing a trade-off between resistance and toxicity in *tetA* expression^36^. Ultimately, tetracycline resistance depends on the interaction between these two inducible mechanisms, which differ in costs/benefits and drug specificities. Resistance can be increased by acquiring mutations in either one.

**Figure 1.**
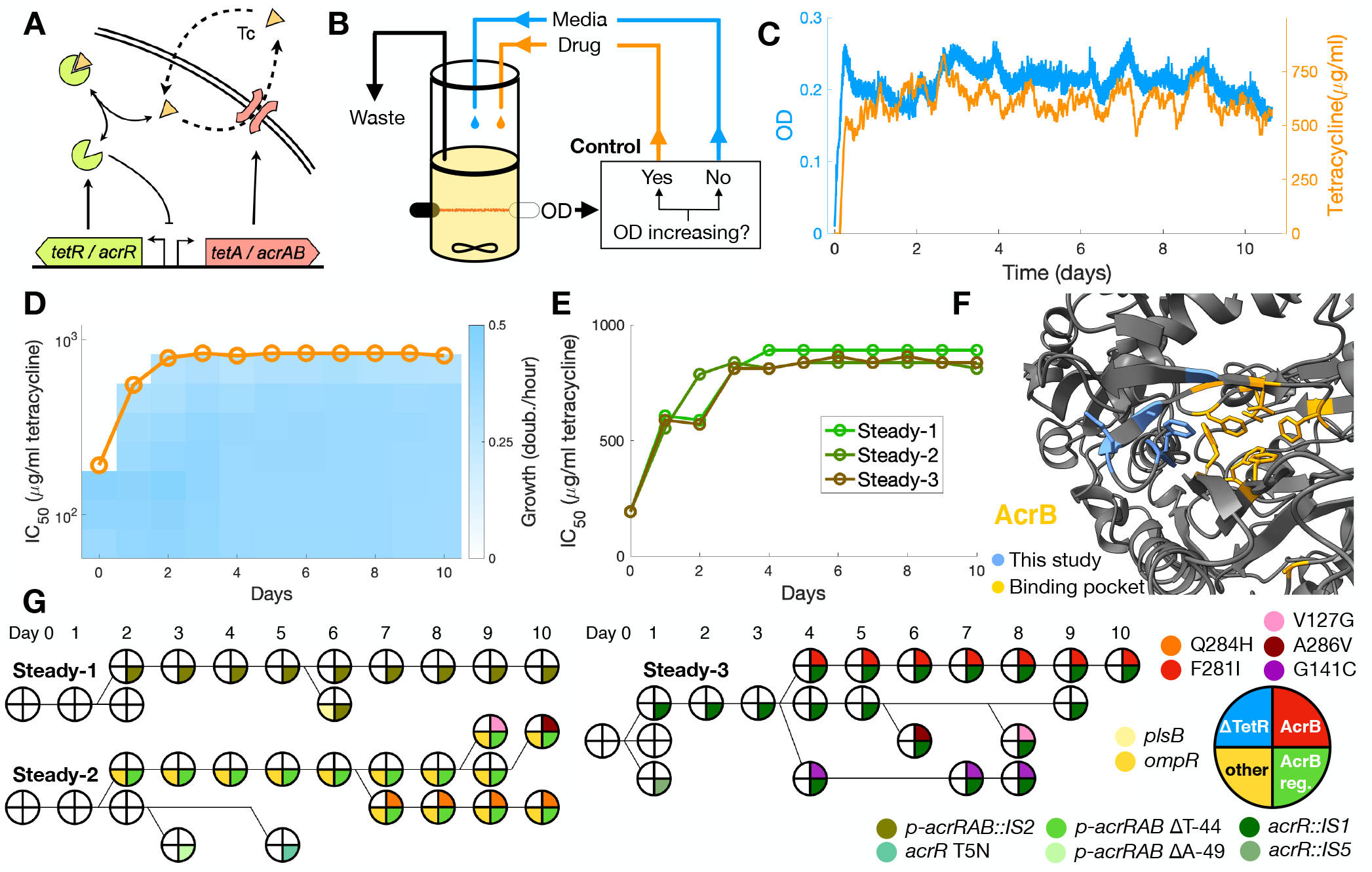
Steady drug regimen favors mutations in the AcrB binding pocket. **A**. *tet* and *acr* resistance mechanisms: tetracycline (Tc), a translation inhibitor, diffuses across the cell membrane and binds transcription repressors TetR and AcrR, which become inactive and release expression of efflux pumps TetA and AcrAB, which export tetracycline out of the cell. While *tet* resistance is specific for tetracycline, *acr* is not. **B**. Continuous culture setup with media and drug control for the Steady environment. If OD increased over the previous half hour, a fixed drug dose is added to the culture. **C**. Progression of OD and drug concentration for population Steady-2 over the course of the experiment. **D**. Growth rates measured in samples of population Steady-2 from each day of the experiment along a gradient of tetracycline. Circles indicate the drug concentrations at which growth is reduced to half (IC_50_ concentration). **E**. Increase in resistance for each Steady population, measured by the IC_50_ concentration. **F**. Locations of AcrB mutations detected in evolved populations within the protein structure. We found several mutations around the drug binding site. **G**. Cladograms of isolates from each Steady population, showing the presence of high-abundance mutations. 5 isolates were picked from each population in each day, and presence of select mutations was confirmed by sequencing. Phylogenetic relationships were inferred from the co-occurrence of each mutation in each isolate.

Here, we use a system of automated continuous cultures to experimentally evolve tetracycline resistant *E. coli* populations under different regimens of drug administration. We compare evolution under a steady drug environment, where the drug concentration changes gradually, with a dynamic environment where the population is periodically subjected to sudden exposures to high drug concentrations.

## Results

We have developed a device for microbial evolution in controlled dynamic environments, based on the morbidostat design^21^, which propagates continuously growing cultures while automatically adjusting media input and antibiotic concentration to impose specific regimens of drug administration. The device allows the maintenance of cultures at a set density by measuring turbidity (optical density) in real time and externally adjusting media and drug delivery via a control algorithm. We used our setup to impose both a “Steady_*i*_ and a “Dynamic_*i*_ environment to the evolving populations, implemented by different control algorithms.

### Steady environment favors mutations adapting AcrB to tetracycline

Most experimental evolution studies of antibiotic resistance have focused on the acquisition of resistance by sensitive strains, while less attention has been given to the evolution of strains already carrying dedicated mechanisms of resistance^37^. Previous evolution studies with a sensitive *E. coli* strain lacking the *tet* operon resulted in only a 10-fold increase resistance to tetracycline, with most adaptive mutations found in genes coding for membrane proteins, including the *acr* operon, or factors affecting transcription and translation^21^. For comparison, strains carrying the *tet* operon (*tet+*) achieve much higher resistance, with a minimum inhibitory concentration (MIC) 100-fold larger than sensitive strains^7^. This raised the question of if the evolution of *tet+* strains would target the *tet* operon itself. If the native *tet* operon could be further optimized for higher doses of tetracycline, mutations in the *tet* genomic locus were likely to have large effects on resistance. Otherwise, if the *tet* operon were already highly optimized, adaptive mutations would be expected elsewhere, either providing resistance through other means or compensating for fitness costs associated with high TetA expression^38–40^.

To determine the evolutionary trajectory of *tet*+ strains to increased tetracycline resistance, we evolved 3 populations (Std-1, Std-2 and Std-3) under a “Steady_*i*_ regimen following the morbidostat design (Fig 1BC, Fig S1, methods). In this regimen, the drug is always present, with levels gradually increasing as the population evolves resistance. Populations were sampled daily throughout the experiment. To measure the increase in drug resistance in each population, we inoculated a sample from each day in a gradient of tetracycline, measuring the drug concentration that reduces growth by half relative to growth in the absence of drug (IC_50_). By the third day, all 3 Steady populations had developed high levels of resistance (Fig 1DE, Fig S2). Next, we sent genomic DNA from each final culture for whole genome sequencing to identify adaptive mutations.

All three Steady cultures evolved genetically diverse populations (Table 1) with significantly increased resistance in comparison to the wild type. We did not find any mutations in the *tet* operon locus, suggesting that TetA is already highly optimized for tetracycline efflux. Instead, we observed a predominance of mutations enhancing other mechanisms of resistance, particularly the *acr* operon. All three populations acquired mutations upregulating AcrB, either by disrupting repressor AcrR or the *acrRAB* promoter region. Additionally, two populations acquired mutations in the drug binding pocket of the AcrB protein itself^34,35^, presumably increasing the affinity of AcrB to tetracycline (Fig 1F).

**Table 1.** Relevant mutations found in evolved Steady populations. Mutations shown are present above a 0.15 frequency and are not detected in the ancestral WT population.*Found in very low abundance in the ancestral WT population.

| Population | Gene | Function | Frequency |
| --- | --- | --- | --- |
| Std-1 | *p-acrRAB::IS2* | Multidrug efflux pump | 0.86 |
| Std-1 | *plsB* | Phospholipid biosynthesis, multidrug tolerance | 0.63 |
| Std-1 | *insB1* | Insertion element IS1 protein | 0.15 |
| Std-1 | *del cheA/*<br>*motAB/flhCD\** | Chemotaxis, motility | 1 |
| Std-2 | *p-acrRAB* | Multidrug efflux pump promoter | 0.99 |
| Std-2 | *ompR* (R15S) | Porin regulator, osmotic stress | 0.97 |
| Std-2 | *yhdP* | Phospholipid transport factor | 0.37 |
| Std-2 | *acrB* (Q284H) | Multidrug efflux pump | 0.36 |
| Std-3 | *acrR::IS1* | AcrB multidrug efflux pump repressor | 0.78 |
| Std-3 | *acrB* (F281I)* | Multidrug efflux pump | 0.75 |
| Std-3 | *p-rrsG* (2729370) | 16S ribosomal RNA | 0.27 |
| Std-3 | *p-rrsG* (2729333) | 16S ribosomal RNA | 0.24 |
| Std-3 | *del cheA/*<br>*motAB/flhCD\** | Chemotaxis, motility | 1 |

Therefore, in Steady populations resistance is increased by optimizing the *acr* operon for tetracycline efflux, complementing the resistance provided by TetA.

In addition to mutations in the *acr* operon, we also observed other mutations that potentially affect membrane permeability to tetracycline, such as in porin regulator OmpR and in PlsB and YhdP, involved in phospholipid synthesis and transport^41,42^. In all evolved populations, there were several low-abundance mutations related to transposable elements (discussed below). Interestingly, population Std-3 showed two mutations upstream of 16S subunit ribosomal RNA *rrsG*, the target of tetracycline, which is possibly upregulated in response to the drug. Interestingly, a large deletion of chemotaxis and motility genes that was already detected in low abundance in the ancestral WT population became fixed in populations Std-1 and Std-3. We speculate that this deletion of several motor proteins involved in motility might alleviate the depletion of the membrane potential caused by constant TetA expression.

Next, we analyzed the emergence of the most abundant mutations in the evolved populations: *plsB* (Std-1), *acrB* (Std-2 and Std-3) and *ompR* (Std-2 and Std-3). We picked 5 isolates from each population for each day of the experiment and verified the presence of mutations in each gene by Sanger sequencing. We then determined the order and co-occurrence of mutations to infer the competing lineages in each population over the course of the experiment (Fig 1G). Surprisingly, we found several competing mutations in the *acr* operon that were not detected in the final populations. Mutations in the *acrB* gene itself were always preceded by mutations disrupting AcrB repression, suggesting that tetracycline is not an effective inducer of the *acr* operon, and upregulation of *acrAB* is necessary to potentialize further mutations in *acrB*. Of notice, population Std-1achieved upregulation of *acrAB* through an *IS2* transposon insertion within its promoter region, with no further mutations anywhere in the *acr* operon. The *IS2* insertion sequence is known to introduce a strong promoter that results in upregulation of adjacent genes^43^. Overall, these results indicate that the evolution of non-optimal resistance mechanisms in steady drug regimens proceeds first by upregulation of the resistance protein, followed by the optimization of the resistance proteins themselves.

### Dynamic environment favors loss of *tet* regulation

Experimental evolution studies of antibiotic resistance have typically been conducted under steadily increasing antibiotic concentrations, which does not account for the dynamic nature of most natural habitats such as clinical settings^44,45^. In these environments, while drug exposures can be relatively rare, they happen suddenly and at high concentrations, requiring regulated mechanisms of resistance to deliver fast responses. At first thought, evolution in such highly dynamic environments should lead to faster induction of resistance. However, faster responses might depend on rare mutations that are not likely to occur under the strict evolutionary bottlenecks imposed by sudden exposures to high drug doses. Therefore, it was unclear if faster regulation of the *tet* operon would be a likely outcome of evolution in such environments.

To determine the role of dynamics in the evolution of drug responses, we evolved three resistant populations (Dyn-1, Dyn-2 and Dyn-3) under a “Dynamic_*i*_ drug environment, implemented by a daily single large dose of tetracycline followed by growth in fresh medium (Fig 2A-C, Fig S1, methods). By adjusting the drug dose based on the previous recovery time of the population, we maintain the evolutionary pressure for faster response times, while the subsequent growth in fresh medium provides a fitness advantage for cells that grow efficiently in the absence of drug. A regimen of daily administration of the drug was chosen to provide enough time for growth in drug-free conditions and to reflect typical timescales of antibiotic treatments. Next, we measured the increase in drug resistance in each population throughout the experiment. Overall, all three Dynamic populations increased resistance more slowly in comparison to the Steady populations, as measured by the IC_50_ (Fig 2DE, Fig S2). These results support the notion that strict evolutionary bottlenecks imposed by the Dynamic environment can select mutants that are less fit in comparison to the Steady environment.

**Figure 2.**
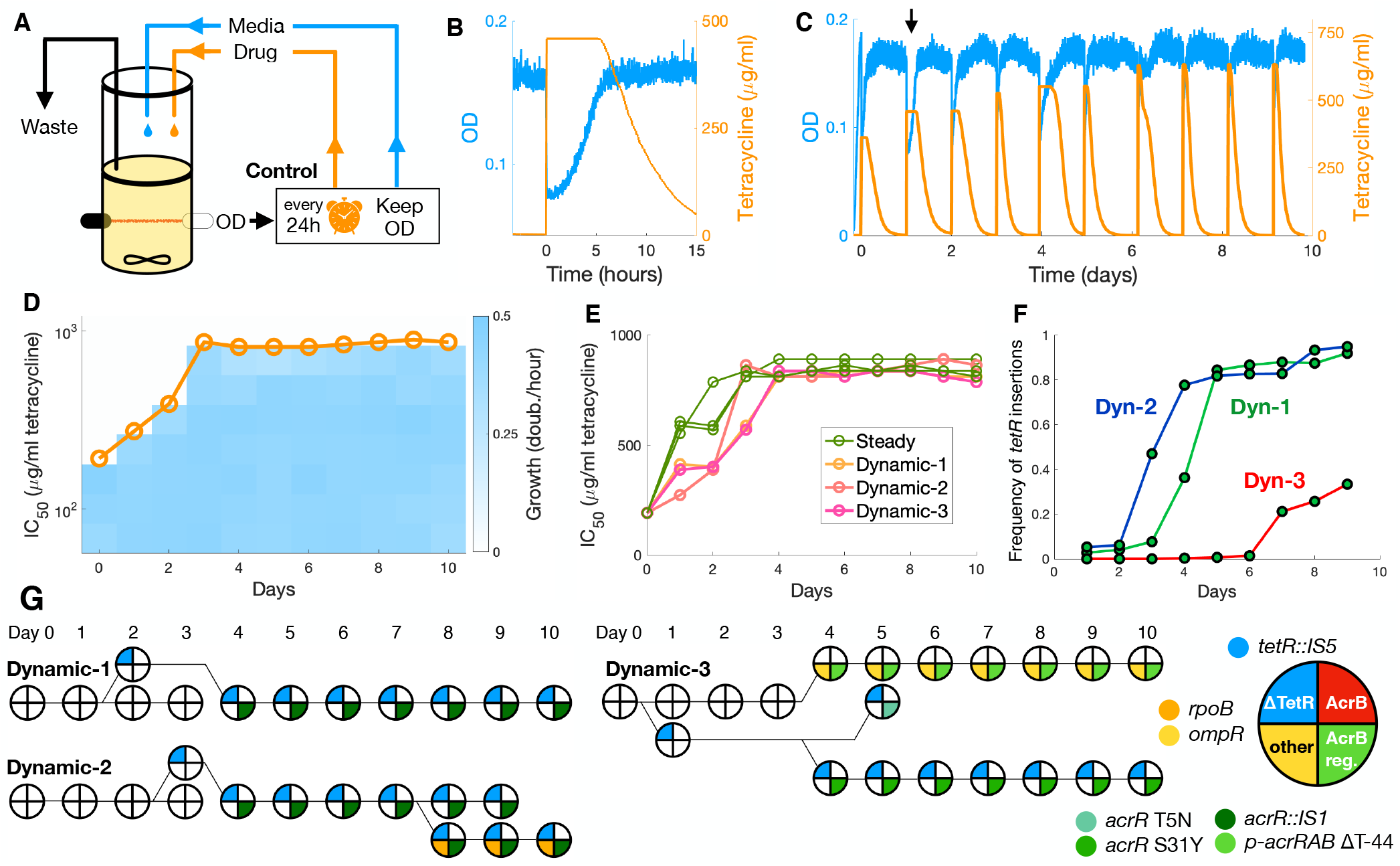
Dynamic drug regimen favors transposon insertions in the *tetR* gene. **A**. Continuous culture setup with media and drug control for the Dynamic environment. Cultures are suddenly exposed to a large drug dose every 24 hours but are otherwise maintained at a target OD by dilution with fresh medium. **B**. Cultures are exposed to tetracycline by the addition of a large volume of medium containing the drug, which reduces OD to half. Dilution with fresh medium resumes when the population doubles to again reach the target OD. **C**. Progression of OD and drug concentration for population Dynamic-2 over the course of the experiment. Arrow indicates the exposure depicted in (B). **D**. Growth rates measured in samples of population Dynamic-2 from each day of the experiment along a gradient of tetracycline. Circles indicate drug concentrations at which growth is reduced to half (IC_50_ concentration). **E**. Increase in resistance for each population, measured by the IC_50_ concentration. Acquisition of resistance is slower in Dynamic populations. **F**. Prevalence of transposon insertions in *tetR* over time in each Dynamic population. Such mutations were the first to be observed in all Dynamic populations. **G**. Cladograms of isolates from each Dynamic population, showing the presence of high-abundance mutations. 5 isolates were picked from each population in each day, and presence of select mutations was confirmed by sequencing. Phylogenetic relationships were inferred from the co-occurrence of each mutation in each isolate.

The Dynamic regimen produced even higher genetic diversity than the Steady regimen (Table 2). Unexpectedly, instead of evolving fast on/off switching of the drug response, all Dynamic populations independently acquired *IS5* transposon insertions in different loci inside the *tetR* gene, resulting in loss of TetR function. We verified the presence of these insertions in each population throughout the experiment by PCR and found that *IS5* insertions in *tetR* rapidly emerged and reached fixation in populations Dyn-1 and Dyn-2 (Fig 2F, Fig S3). In population Dyn-3, *tetR* insertion only rose in frequency in the last few days but did not reach fixation. Such insertions abolish regulation of efflux pump TetA, which becomes permanently de-repressed. Still, TetA levels are expected to be similar between *tetR* mutants and WT cells when fully induced in the presence of the drug. These results suggest that constitutive expression of TetA improves resistance by bypassing the induction of the response, and therefore we hypothesized that preemptive expression of resistance could shorten population-level recovery to sudden drug exposures despite the known fitness costs of TetA overexpression.

**Table 2.**
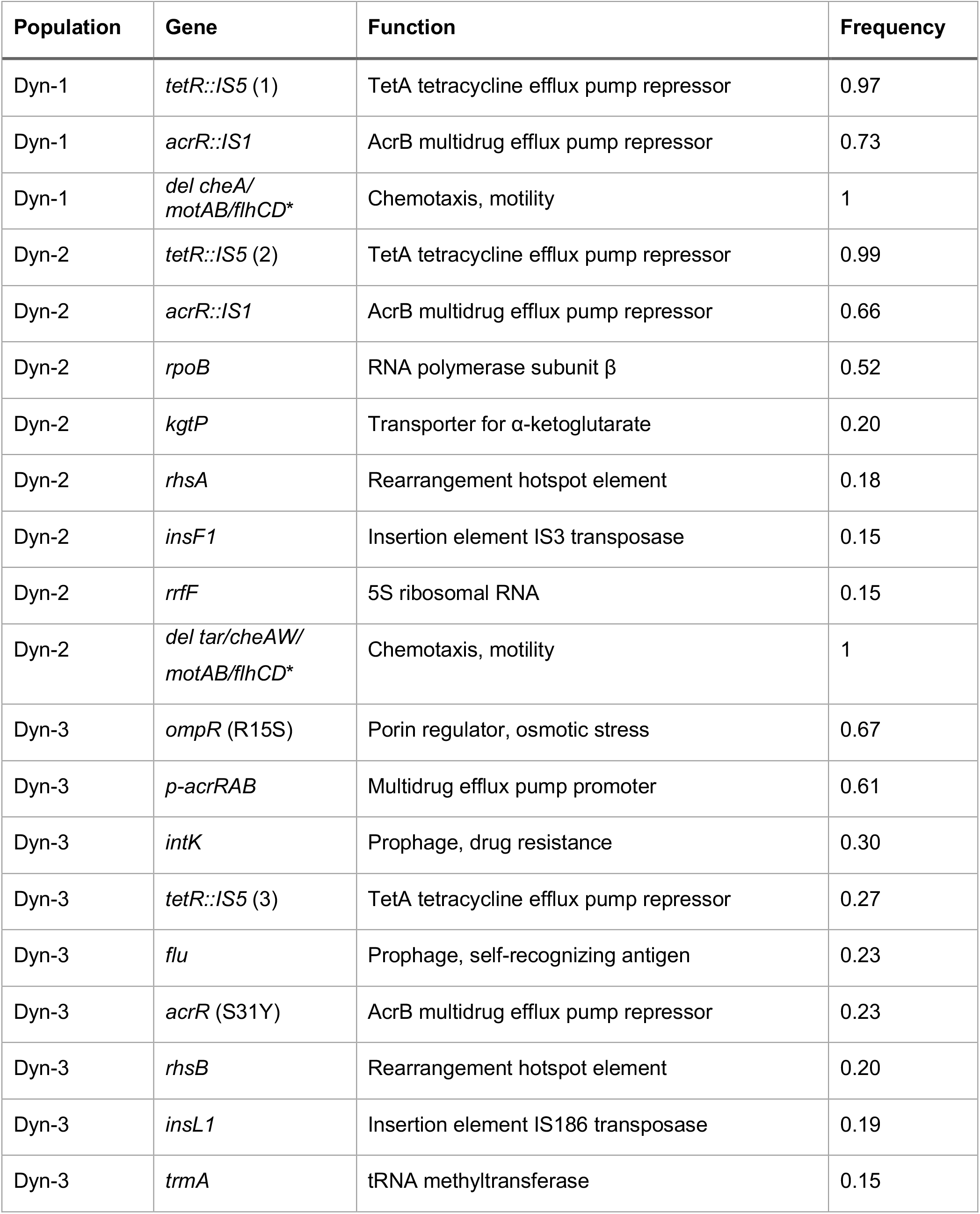
Relevant mutations found in evolved Dynamic populations. Mutations shown are present above a 0.15 frequency and are not detected in the ancestral WT population.*Found in very low abundance in the ancestral WT population.

In all Dynamic populations, we also observed a high abundance of *acr* regulatory mutations. However, we did not find any mutations in the *acrB* gene. Populations Dyn-1 and Dyn-2 both showed *IS1* transposon insertions in repressor *acrR*, and population Dyn-3 showed both mutations in the *acrRAB* promoter and in *acrR* itself. Therefore, upregulation of AcrB also increases fitness in the Dynamic environment. Again, we observed other prevalent mutations that potentially reduce membrane permeability to tetracycline or compensate for its effects, such as *ompR*, RNA polymerase subunit *rpoB* and the *kgtP* transporter, as well as several mutations related to transposable elements. In populations Dyn-1 and Dyn-2, we also found the fixation of the large deletion of chemotaxis and motility genes that might help compensate for TetA overexpression.

Again, we picked 5 isolates from each population for each day of the experiment and verified the presence of the most abundant mutations: *acrR* (Dyn-1, Dyn-2 and Dyn-3), *rpoB* (Dyn-2) and *ompR* (Dyn-3). We then determined co-occurrence of mutations within the same lineages to infer the competing lineages in each population (Fig 2G). In all populations, *tetR* insertions preceded any other mutations and were inevitably followed by mutations upregulating the *acr* operon. However, while *tetR* insertions were quickly fixed in populations Dyn-1 and Dyn-2, population Dyn-3 showed an interesting case of clonal interference. Although a *tetR* insertion was detected early, by day four it was competing with another lineage carrying mutations in *ompR* and in the *acRAB* promoter. The lineage with the *tetR* insertion then developed two competing *acr* regulatory mutations before rising to ∼30% abundance in the last 4 days of the experiment. Therefore, mutations outside of the *tet* operon can also significantly increase fitness in dynamic environments and interfere with the emergence of *tetR* mutants. Next, we moved to study the dynamics of the response in the evolved populations and in relevant isolates.

### Mutations optimizing *acrB* provide highest resistance and shortest recoveries

To measure antibiotic resistance in the evolved populations during abrupt changes in drug concentration, we developed a liquid-culture assay based on the dynamics of population-level growth during drug responses^46,47^ (Fig. 3A). Sudden exposures to high drug concentrations cause the growth of liquid cultures to pause temporarily, as a large subpopulation of cells is arrested^7^. Growth is then resumed when the growing subpopulation of surviving cells surpasses this background of arrested cells. We then quantify resistance using two components: “steady-state resistance_*i*_, which measures the capacity for bulk growth in the presence of drug (similar to IC_50_), and “dynamical resistance_*i*_, which measures the speed of population-level recovery as the delay in the time it takes the population to achieve one doubling following drug exposure (methods, Fig S4-7). Both the speed of recovery and the ensuing steady-state growth rate decline as a function of the drug concentration used in the exposure.

**Figure 3.**
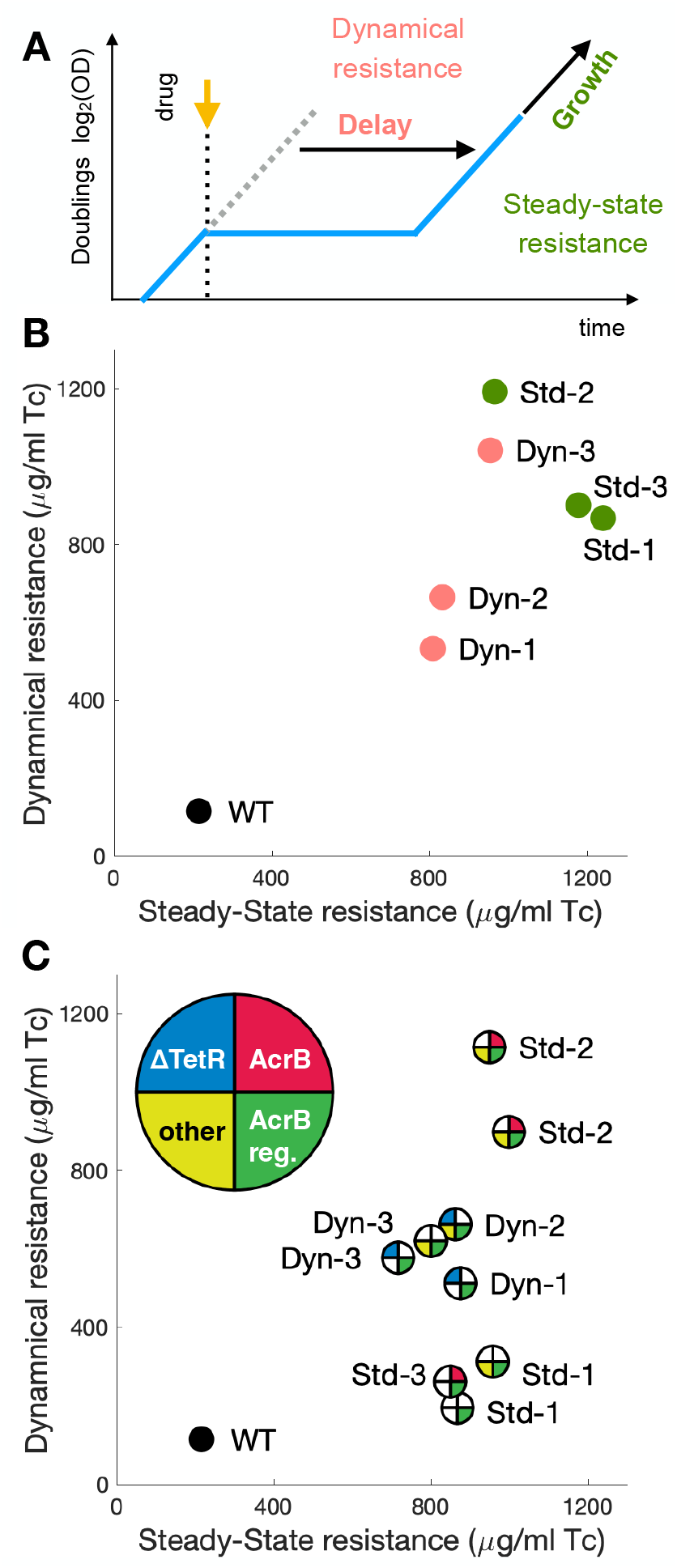
Mutations optimizing *acrB* provide highest resistance and shortest recoveries. **A**. In experiments where liquid cultures are exposed to a step increase in drug concentration, we calculate *Dynamical Resistance* to measure the population-level capacity for quickly recovering growth following exposure, using the delay in the time taken to reach one doubling that is introduced by the addition of drug during mid-log phase. We calculate *Steady-state Resistance* to measure the capacity for steady-state growth in the presence of drug, using the maximum growth rate reached after exposure. **B**. Dynamical and Steady-state resistances for the final evolved populations. Populations Dyn-1 and Dyn-2, where *tetR* transposon insertions were fixed, performed worse in both measures. **C**. Dynamical and Steady-state resistances for select isolates. *tetR* transposon mutants showed high dynamical resistance, but mutants with mutations optimizing AcrB for tetracycline efflux performed best.

While all populations evolved similar levels of steady-state resistance, we found significant differences in dynamical resistance. Surprisingly, populations evolved in the Dynamic regimen showed lower dynamical resistance than the populations evolved in the Steady regimen (Fig. 3B, Fig S5). Therefore, mutations acquired under constant drug pressure can also shorten population recovery following sudden drug exposures. We also note that all evolved populations grow slightly faster in the presence of moderate concentrations of tetracycline than in its absence (Fig S5). Since the evolved populations harbor several mutants, we also assessed resistance in a collection of representative isolates (Fig. 3C, Table S2, Fig S6). We found that while all isolates from the Dynamic regimen showed increased dynamical resistance compared to the WT, the highest dynamical resistance was measured in isolates from Steady population Std-2 combining mutations in *acrB* and *ompR*. Therefore, mutations improving the efficiency of tetracycline efflux are likely to result in both faster recovery from drug exposures and faster steady-state growth at high drug concentrations. Meanwhile, regulatory mutations that shorten recovery from exposure, such as *tetR* transposon insertions, do not necessarily improve steady-state growth at high drug levels.

### Sudden exposures to high drug doses overwhelm regulated responses

To understand the fitness advantage provided by loss of *tetR* repression, we picked isolates from the same sample of early Dynamic populations where *tetR* insertions first emerged. We then compared how isolates with and without *tetR* insertions respond to sudden exposures to tetracycline (Fig. 4A). Liquid cultures of isolates with *tetR* insertions recovered growth much earlier following exposure, despite little change in the steady-state growth rate once growth was recovered. Therefore, abolishing TetR repression of TetA increases cell survival to the initial exposure and recovery of population growth, but steady-state growth under full TetA induction is still similar between WT and *tetR* mutants.

**Figure 4.**
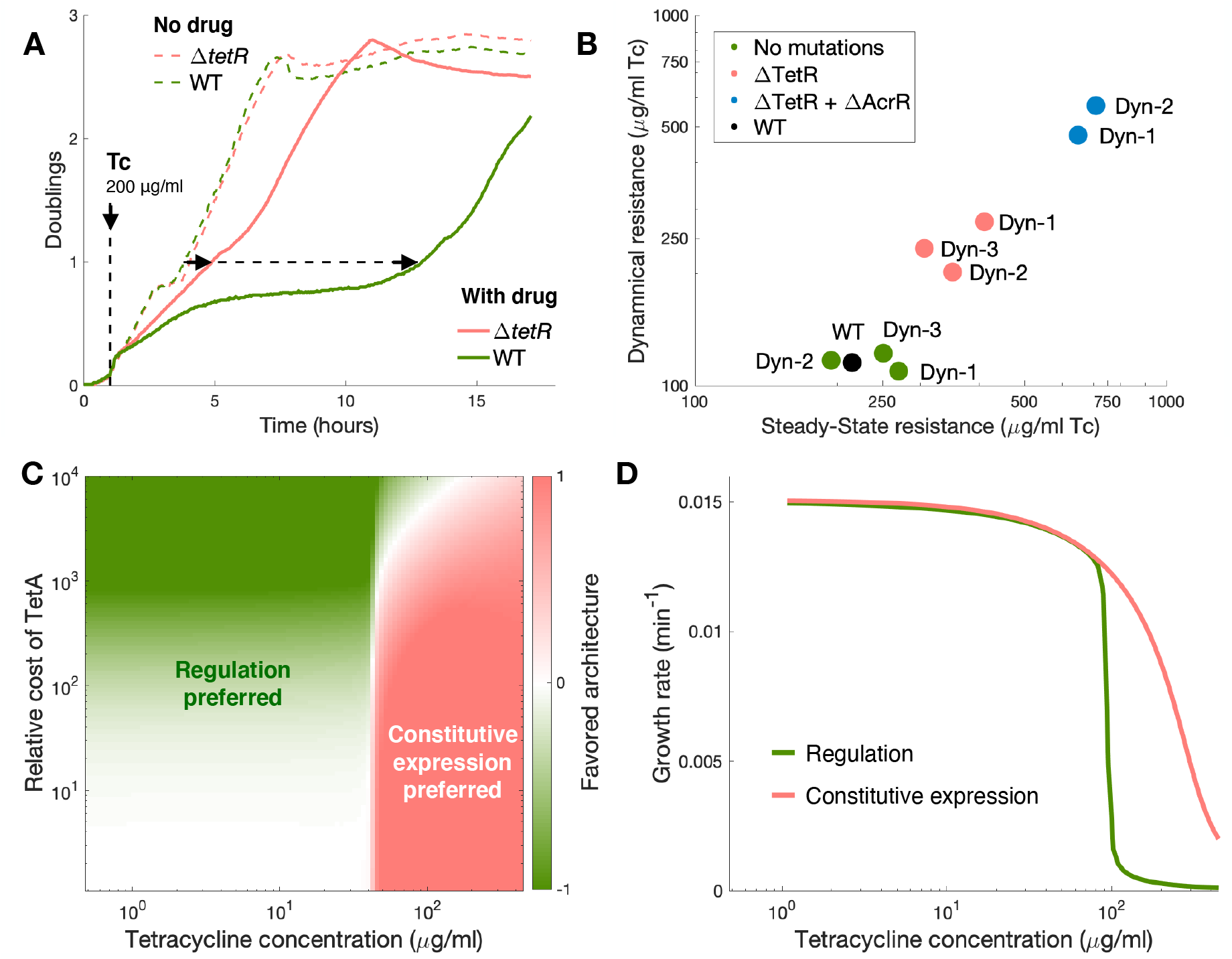
Sudden exposures to high drug doses overwhelm regulated responses. **A**. Growth curves of isolates picked from the same day, one with a transposon insertion in the *tetR* gene and one without. The vertical dashed line indicates addition of 200 μg/ml tetracycline. The corresponding growth curves under no drug are also included in dashed lines for comparison. The delays for each culture to reach one doubling following drug exposure is indicated by dashed arrows. Loss of TetR function reduces the recovery time significantly. **B**. Dynamical and Steady-state resistances for isolates with and without *tetR* transposon insertions, and also with further regulatory mutations in the *acr* operon. Loss of TetR function provides significant increase in resistance, but further upregulation of the *acr* operon increases resistance even further. **C**. We simulate drug responses using a mathematical model of the drug response. We simulated the system with and without TetR repression, during exposures to a range of tetracycline concentrations. To compare the two regulatory strategies, we calculated costs associated with the decrease in cell growth and with the burden of unnecessary TetA production, considering scenarios where constitutive TetA expression has different relative contributions to the overall cost of the response. Regardless of the relative cost of TetA, constitutive expression is preferred during exposures to high drug doses. **D**. Simulations showing final steady-state growth following exposures to different drug concentrations, for both regulated and constitutive expression of resistance. There is a significant window of selection where constitutive expression of TetA allows for the maintenance of cell growth during drug exposures, while TetR-regulated TetA expression fails to induce a response.

Once *tetR* mutations are acquired, upregulation of *acrB* provides further increase of both recovery speed and steady-state growth. We measured dynamical and steady-state resistance in WT, *tetR* mutants and in *tetR* + *acrR* mutants (Fig. 4B, Fig S7). *tetR* insertions increased dynamical resistance substantially, while providing only a slight increase in steady-state resistance. Therefore, since WT and *tetR* mutants showed similar steady-state growth rates for all drug concentrations where they are both viable (Fig S7), the selective advantage of *tetR* mutants likely results from WT cells not being able to fully induce TetA expression in time. Contrastingly, further acquisition of *acrR* mutations by *tetR* mutants resulted in much higher gains in both steady-state and dynamical resistance, corroborating the hypothesis that mutations that improve tetracycline efflux increase resistance in any regimen of drug administration.

To understand how sudden drug exposures can select for loss of TetR repression, we developed a mathematical model of the dynamics of drug responses^48–50^ that tracks the accumulation of intracellular tetracycline and expression of both TetA and TetR. This model includes drug effects on growth and gene expression^51,52^, and loss of TetR repression is modeled by simply reducing TetR expression to zero (methods, SI). For both the presence and absence of TetR regulation, we simulated the model in a range of external tetracycline concentrations (Fig. 4CD). We then compared the two regulatory strategies by calculating costs associated with the decrease in cell growth and the burden of unnecessary TetA production in each case. Since the fitness cost of TetA is difficult to determine, we considered different values for the relative contribution of TetA expression to the overall cost of the response. Calculating the overall cost of the drug response for each regulatory strategy, we found that, regardless of the burden associated with TetA expression, regulated TetA expression is optimal during exposures to lower drug concentrations and constitutive TetA expression is optimal at higher drug concentrations. Therefore, since exposure to high drug concentrations can result in drug influx outpacing expression of resistance, constitutive expression is preferred despite its potential costs.

### Mobile elements as a general strategy of mutating regulatory pathways

Both the Steady and Dynamic environments resulted in an abundance of regulatory mutations, and many of these were implemented by the insertion of transposable elements. Crucially, all Dynamic populations evolved independent *IS5* transposon insertions inside the *tetR* gene, making TetA expression constitutive. All six populations from both environments evolved upregulation of *acrB*, and four of these mutations were also mediated by transposon insertions. Overall, the highest frequency mutations included several examples of disrupted transcriptional repressors and their binding sites, with about half of these regulatory mutations mediated by transposon insertions. Apart from disrupting regulation, transposon insertions also introduced an additional promoter for *acrB* and mediated the large deletion of chemotaxis and motility genes. Populations in both environments also exhibited a wealth of low-frequency mutations in genes related to mobile genetic elements^53,54^ (Tables 1-2). Comparatively fewer mutations were observed in non-regulatory genes (*acrB, rpoB* and *plsB*), and these mutations only appeared later in the experiment in lineages that already possessed regulatory mutations. Thus, these results support the notion that microbial evolution in new environments proceeds first through regulatory mutations, which are more accessible than the low probability point mutations necessary to reach optimal phenotypes. Mobile genetic elements then provide a fast mechanism for upregulating useful genes, either by disrupting repressors or inserting new promoters, or also deleting harmful genes.

## Discussion

Performing experimental evolution using carefully controlled continuous cultures, we studied the role of the dynamics of drug delivery in the evolution of antibiotic responses. We found that steady environments where drug concentrations change only gradually led to the refinement of the AcrB efflux pump through point mutations that optimize it for tetracycline efflux. Meanwhile, dynamic environments with sudden exposures to large drug concentrations did not result in such refinement of protein function, but instead led to the abolishment of regulation of the *tet* operon, the main mechanism of tetracycline resistance. Such mutation was unexpected in a changing environment, since loss of regulation had been previously understood to be a long-term outcome of evolution in constant environments^55^. These experiments show that regulation is not only important for the parsimonious use of cellular programs that are only needed occasionally, but also needs to guarantee a timely activation of cell defenses in hostile environments, while gene expression is still possible.

Regardless of environment, evolution proceeded first through regulatory mutations^56–58^, with a high prevalence of transposon insertions. Previous work shows that exposure to low levels of tetracycline results in high levels of mobile element activity via the sensing of cellular stress^59–61^. Evolution under high drug levels requires quick adaptation, and transposon insertions provide an accessible mechanism for genome remodeling. Transposon insertions also offer the advantage of being reversible^62^. The overwhelming presence of transposon insertions rewiring the regulation of resistance genes, as well as mutations in genes regulating and operating transposable elements, indicates an important role of this mechanism in the short-term evolution of microbes in challenging environments, which is often missed in whole-genome sequencing analysis.

Sudden drug exposures are likely to play a role in the evolution of pathogens in clinical environments. Most antibiotic treatments are well designed to reach and maintain high drug concentrations, but short courses such as intravenous or inhalation result in sharp increases of drug bioavailability at the site of infections^63^. Indeed, loss-of-function mutations in transcription repressors of resistance genes are often found in clinical isolates^64,65^. Our results suggest that these mutations can serve both the purposes of bypassing the induction of drug responses and of de-repressing resistance mechanisms that are not sufficiently activated by the drug. Constitutive expression of resistance would be even more beneficial for resistance mechanisms that are induced only by downstream effects of the drug and respond much more slowly than the *tet* operon^66^.

This work raises interesting questions about how the dynamics of the environment shapes evolution, such as whether dynamic regimens with lower drug concentrations might ease evolutionary bottlenecks and ultimately select for faster responses, or whether longer periods in the absence of drug might reduce the benefits of constitutive expression of resistance. Dynamic environments could also promote the coexistence of multiple mutants or species that are optimized for different drug levels, shaping the composition and evolution of complex microbial communities^67^.

## Methods

### Media, drugs, and strains

All experiments were conducted in M63 minimal medium (2g/l (NH4)2SO4, 13.6g/l KH2PO4, 0.5mg/l FeSO4,7H2O) supplemented with 0.2% glucose, 0.01% casamino acids, 1mM MgSO4 and 1.5mM thiamine. Tetracycline and IPTG solutions were freshly made from powder stocks (Sigma) and filter-sterilized before each experiment. As the ancestral strain, we used *E. coli* K-12 strain MG1655 *rph*+ Δ*LacIZYA*, with the native *tet* resistance mechanism from the Tn*10* transposon integrated in the chromosome at site HKO22 with a pIT3-CH integrating plasmid.

### Setup of continuous cultures

Glass vials were coated with Sigmacote (Sigma) to prevent biofilm growth. Vials, vial heads, tubing, and all connections were autoclaved before assembly. Complete sterilization was ensured by running 10% bleach, 70% ethanol, and sterile water consecutively through all tubing. Experiments were carried out at 37 °C in sterile M63 media (as described above). Drug media consisted of M63 media with tetracycline added to 1000 µg/ml, mixed until completely dissolved in solution, and filter sterilized into the container used for the experiment. Vials were filled with 15 ml of M63 and used to blank OD measurements.

### Continuous culture experiments

The experiment consisted of two experimental groups (Dynamic and Steady); each group was composed of three replicate vials. Samples were taken daily and stored as glycerol stocks in a 96 well plate at -80 °C. Vials were switched daily to prevent biofilm growth.

### Steady regimen

The steady group was run as a morbidostat. We used 15 ml of cell cultures at a fixed dilution rate of 0.35 h^-1^, with media added every 5 minutes, which corresponds to a growth rate of 0.5 doublings per hour and an OD (optical density) of 0.15 for the ancestral strain in the absence of drug. Tetracycline was automatically added to the cultures in increments of 4.4 µg/ml in every cycle where the population showed net growth over the previous 30 minutes. This regimen initially resulted in cell densities between 0.2 and 0.3, while keeping a constant selective pressure resulting from drug inhibition of growth. After 3 days, all 3 populations were growing stably with OD around 0.3 and drug concentrations above 500 µg/ml, which is ∼5-fold higher than the MIC of the ancestral WT strain.

### Dynamic regimen

The dynamic group was run as a modified turbidostat. We kept 15 ml of cell culture at a fixed target OD of 0.15 by the controlled dilution of the culture with drug-free media, delivered every 5 minutes. At 24 hour intervals, cultures were exposed to a single large dose of tetracycline by the addition of 15 ml of medium with drug, which also caused the population to be diluted by half. We then waited for the culture to recover to the target OD without dilution. Following recovery, controlled addition of fresh medium resumed, and the drug concentration gradually decreased to zero (Fig 2B). We started with an exposure to 150 µg/ml of tetracycline in the first day, which was shown to cause a ∼5 hour delay in recovery from drug exposure in the ancestral WT population. The drug concentrations for subsequent exposures were adjusted by a factor of 5/ *t*_*r*_, where *t*_*r*_ Is the recovery time in hours, which resulted in an increase in drug concentration if the recovery time was shorter than five hours. If a culture did not fully recover within 24 hours, no drug was added in that cycle.

All three populations quickly recovered from the first exposure and were followed by significant increases in drug dose for the exposures in the following days. Recovery times were longer and more variable within the first few days, but eventually stabilized at around 45 to 60 minutes by the end of the experiment (Fig 2C, SI). In general, populations recovered quickly and showed fast growth following drug exposures, which resulted in fast dilution rates and ensured that the drug was removed from the culture within 10 hours.

### Genomic sequencing of evolved populations

Samples were collected daily during the experiment. On completion, gDNA from each final culture was extracted and sent for whole genome sequencing, together with the ancestral WT population (MG1655 *tet+*), to identify adaptive mutations by aligning WGS reads to a reference genome. We also sequenced a culture evolved in a turbidostat in the absence of drug to identify possible mutations that are adaptive to growth under our experimental conditions, but not related to drug resistance. We did not find any significant such mutations.

Glycerol stocks from the final evolved populations were inoculated 1:100 in 5 ml LB and grown for five hours. Samples were pelleted and DNA was extracted using DNEasy Kits (Qiagen). Whole genome sequencing, producing a median of 2.3 million, 2×151bp, paired-end, reads was performed by the Genomics Shared Resource, and sequencing analysis was carried out by the Genomic Data Science Core at Dartmouth College.

### Analysis of whole genome sequencing

Raw reads were trimmed using Trimmomatic v.0.39^68^ and the following parameters: “ILLUMINACLIP:TruSeq3-PE-2.fa_wpoly.fa:2:30:10:2:keepBothReads LEADING:3 TRAILING:3 MINLEN:36”. Reads were subsequently aligned to the *E. coli* K-12 U00096.2 reference genome using BWA v.0.7.17^69^. Single nucleotide variants and small insertions or deletions were called using FreeBayes v.1.3.4^70^ and annotated using SnpEff v5.0e^71^. An augmented genome with both the U00096.2 reference genome and the integrated plasmid with the *tet* resistance mechanism (above) was used to determine transposon insertion sites within the plasmid. Briefly, sites containing discordant paired end reads were collected whenever one end of the read aligned to the plasmid and one end aligned to the *E. coli* genome. Reads aligned to these collected sites were then tallied as either matching the genome reference or containing a breakpoint, using a custom python script available in our GitHub page.

### Genotyping of isolates

The presence of selected SNPs in isolates was confirmed via Sanger sequencing. Glycerol stock samples for each population for each day were inoculated 1:100 in LB and grown for 2 hours, then streaked on LB agar plates and grown at 37 °C overnight. 10 individual colonies from each sample were picked, selected genomic regions were PCR amplified and cleaned (Qiagen) and sent for sequencing to the Dartmouth College Genomics and Molecular Biology Shared Resources Core. Isolates known to have SNPs were preserved as glycerol stocks at -80 °C. For samples with transposon insertions, the PCR product was assessed by gel electrophoresis to confirm insertions. In *tetR* mutants, we confirmed loss of TetR function by transformation with a plasmid expressing mCherry from the *tet* promoter, which results in high fluorescence levels in the absence of TetR repression.

### Plate reader experiments

Glycerol stock of relevant samples were inoculated into M63 or LB media and grown overnight. Each strain was inoculated 1:1000 in M63 and exposed to a gradient of 12 tetracycline concentrations (100-1000 µg/ml) during early log phase. Experiments were carried out at 37 °C in a Synergy Neo2 plate reader (BioTek). We measured both the time until one doubling was achieved following exposure and the steady-state growth rate after recovery.

### Dynamical and Steady-state resistances

From the plate reader experiments, we extract two complementary measures of resistance: 1) *Dynamical Resistance*, which is the drug concentration that introduces a 2-hour delay in the time to reach one doubling following drug exposure in comparison to the absence of drug, and 2) *Steady-State Resistance*, which is the drug concentration that halves the growth rate of the recovered population (similar to the IC_50_ concentration).

### Mathematical Model

A full description of the model and its parameters is available in the SI, and a complete analysis is available in Stevanovic et al^50^. In short, we model the main biochemical interactions involved in the response as a set of differential equations that describe changes in the concentrations of the intracellular drug (*d*), the efflux pump TetA (*a*), and the repressor protein TetR (*r*):

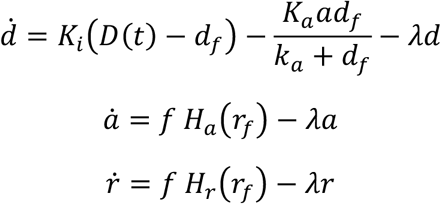

where *K*_*i*_ stands for the import rate, D for extracellular drug concentration, *d*_*f*_ for free intracellular drug, *K*_*a*_ for the catalytic rate constant of TetA, *k*_*a*_ the Michaelis constant, respectively, *r*_*f*_ free repressor, and *H*_*a*_ (*r*_*f*_) and *H*_*r*_(*r*_*f*_)0 the synthesis rates for TetA and TetR that depend on free TetR. The terms *λd, λa* and *λr* represent the dilution due to cell growth of drug, TetA and TetR, where *λ* is the current growth rate. Both the growth rate *λ* and *f*, which allows us to modulate the strength of gene expression according to the proteome partition, depend on nutrient levels and the intracellular drug concentration. To model these dependencies, we used the framework introduced by Scott et al^51^.

### Simulations

We numerically integrate the system of differential equations using the ode113 function in MATLAB, using values of external drug concentration and as an input parameter. The system starts in equilibrium in the absence of drugs, and then the external drug concentration is switched on. The parameter values used in the simulations are summarized in the SI and were either estimated from our experimental data^7^, or obtained from literature^30^. To simulate loss of TetR function, we set *H*_*r*_ *(r*_*f*_) = 0. To calculate the relative fitness of strains with WT regulation vs. constitutive TetA expression, we attributed costs to a decrease in the final steady-state growth rate following drug exposure *c*_*nλ*_ = 1 /*λ*_*final*_ and to the unnecessary expression of TetA in the absence of drugs *c*_*a*_ = *a*_0_ (in the absence of TetR regulation, TetA is fully expressed at all times). Since the relative fitness costs of decrease in resistance and TetA expression are hard to compare, and highly depend on the environment (e.g. frequency and duration of exposures), we introduced a variable weight *w* adjusting the cost of TetA expression, for a total cost of *c* = *w c*_*a*_ + *c*_*λ*_. The fitness advantage of loss of TetR function was then calculated as the relative cost between constitutive TetA expression and WT regulation log (*c*_Δ *TetR*_ /*c*_*wt*_).

## Data Availability

The data that support the findings of this study are openly available. All data and code generated in the study are included in the SI and at our GitHub page: https://github.com/schultz-lab/Evolution-Dynamics. Further inquiries can be directed to the corresponding authors.

## Author Contributions

D.S. and J.C. designed the study. J.C. performed the experiments, and all authors analyzed the data. T.S and J.C. analyzed genomic data. D.S. and H.G. designed the mathematical model. D.S. and J.C wrote the manuscript with input from all authors.

## Acknowledgements

We thank Ben Ross for a detailed reading of our manuscript and many important suggestions. Genomic sequencing was carried out in the Genomics and Molecular Biology Shared Resource at Dartmouth, which is supported by NCI Cancer Center Support Grant 5P30CA023108 and NIH 1S10OD030242 awards. Analysis of sequencing data was performed by the Genomic Data Science Core through the Dartmouth Center for Quantitative Biology with support from NIGMS P20GM130454 and NIH S10OD025235 awards.

## Supplemental Information

## Mathematical Model

A complete analysis of this model is published in Stevanovic et al^50^. We model the main biochemical interactions involved in the response as a set of differential equations that describe changes in the concentrations of the intracellular drug (*d*), the efflux pump TetA (*a*), and the repressor protein TetR (*r*):

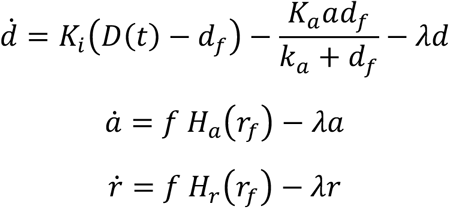

where *K*_*i*_ stands for the import rate, D for extracellular drug concentration, *d*_*f*_ for free intracellular drug, *K*_*a*_ for the catalytic rate constant of TetA, *k*_*a*_ the Michaelis constant, respectively, *r*_*f*_ free repressor, and *H*_*a*_ +*(rf)*and *H*_*r*_ +*(rf)*the synthesis rates for TetA and TetR that depend on free TetR. To simulate loss of TetR function, we set *H*_*r*_+(*r*_*f*_ ) = 0. Since the biochemical binding and unbinding of the substrate to the transcription factor typically happens at a much faster rate than the aforementioned processes, we consider their unbound (free) forms (*d*_*f*_, *r*_*f*_) to be in chemical equilibrium with the bound form [*rd*] with a dissociation constant *K*_/_, such that *r*_*f*_ *d*_*f*_ = [*rd*]. The term *K*_*i*_+(*D* (*t*) − *d*_*f*_)0 represents the diffusion of drug across the membrane into the cell, 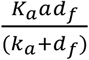the export of drug out of the cell by the efflux pump. The terms *λd, λa* and *λr* represent the dilution due to cell growth of drug, TetA and TetR, respectively, where *λ* is the current growth rate. Both *f*, which allows us to modulate the strength of gene expression, and the growth rate *λ* depend on the nutrient level and the intracellular drug concentration.

To model these dependencies, we used the framework introduced by Scott et al^51^. In this framework *k*_*n*_ and *k*_*t*_ are the nutritional and translational capacity of the cell, respectively. Here, we refer to their base values (full nutrients, no drug) as 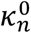 and 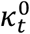, respectively.

Since cultures are kept at lower densities, in regimes where growth is not yet limited by nutrient depletion, we assume the nutritional capacity to remain at 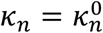. For the translational capacity, we assume inhibition by tetracycline according to 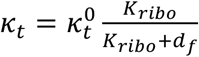, where the fraction represents the fraction of free ribosomes in the cell, and *K*_*ribo*_ is the dissociation constant for tetracycline and the ribosome. Lower *K*_*ribo*_ values correspond to stronger inhibition, and vice versa. The growth rate can then be calculated as 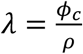.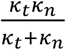,where *ϕ* = 0.48 and *ρ* = 0.76 are universal constants.

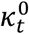 has a universal value of 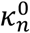 for *E. coli*.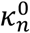 varies according to nutrient quality. We chose 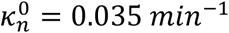 such that 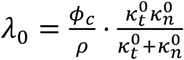 corresponding to a state without tetracycline) yields a maximum growth rate of *λ*_0_= 0.015 *min*^−1^ as observed in our experiments with M63 medium.

We assume that TetA and TetR are part of the P sector of the proteome, which scales as 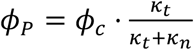 Without regulation of the synthesis rates *H*_*a*_(*r*_*f*_) and *H*_*r*_(*r*_*f*_), we would expect the concentrations of TetA and TetR to scale in the same way. Hence, we define 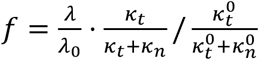, which leads to steady states proportional to 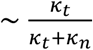 and simplifies to *f* = 1 at full nutrients and no drug.

Parameters were either obtained from literature^30^ or estimated from experiments^7^.

## Model parameters

**Table S1.** Definitions and values of parameters used in the mathematical model.

| Parameter | Value | Definition | Parameter | Value | Definition |
| --- | --- | --- | --- | --- | --- |
| $K_i$ | $0.015 \text{ min}^{-1}$ | drug diffusion influx | $D(t)$ | varies | extracellular drug |
| $K_a$ | $50 \text{ min}^{-1}$ | catalytic rate TetA | $k_a$ | $10 \text{ } \mu\text{M}$ | Michaelis constant TetA |
| $K_d$ | $0.001 \text{ } \mu\text{M}$ | TetR/Tc dissoc. constant | $K_{ribo}$ | $1 \text{ } \mu\text{M}$ | Tc/ribo dissoc. constant |
| $H_a(r_f)$ | $A \frac{r_{0,a}^4}{r_{0,a}^4 + r_f^4}$ | TetA synthesis regulation | $H_r(r_f)$ | $R \frac{r_{0,r}^4}{r_{0,r}^4 + r_f^4}$ | TetR synthesis regulation |
| $A$ | $0.008 \text{ } \mu\text{M min}^{-1}$ | maximal TetA synthesis | $R$ | $0.0003 \text{ } \mu\text{M min}^{-1}$ | maximal TetR synthesis |
| $r_{0a}$ | $0.0001 \text{ } \mu\text{M}$ | dissoc. constant TetA regulation | $r_{0r}$ | $0.000075 \text{ } \mu\text{M}$ | dissoc. constant TetR regulation |
| $\phi_c$ | 0.48 | proteome partition const. | $\rho$ | 0.76 | conversion constant |
| $\kappa_t^0$ | $0.075 \text{ min}^{-1}$ | translational capacity | $\kappa_n^0$ | $0.035 \text{ min}^{-1}$ | nutritional capacity |
| $\lambda_0$ | $0.015 \text{ min}^{-1}$ | max. growth in M63 medium | | | |

**Table S2.** Isolates assessed in Figure 3C. These isolates were picked from final evolved populations and represent the main mutations obtained in this study.

| Population | Gene | Function |
| --- | --- | --- |
| Std-1 | *p-acrRAB::IS2* | Multidrug efflux |
| Std-1 | *p-acrRAB::IS2 + plsB* | Multidrug efflux, phospholipid biosynthesis |
| Std-2 | *p-acrRAB + ompR + acrB (Q284H)* | Multidrug efflux, porin regulator |
| Std-2 | *p-acrRAB + ompR + acrB (A286V)* | Multidrug efflux, porin regulator |
| Std-3 | *acrR::IS1 + acrB (F281I)* | Multidrug efflux, AcrB repressor |
| Dyn-1 | *tetR::IS5 + acrR::IS1* | TetA repressor, AcrB repressor |
| Dyn-2 | *tetR::IS5 + acrR::IS1 + rpoB* | TetA and AcrB repressors, RNAP subunit $\beta$ |
| Dyn-3 | *tetR::IS5 + acrR (S31Y)* | TetA repressor, AcrB repressor |
| Dyn-3 | *p-acrRAB + ompR* | Multidrug efflux, porin regulator |

**Figure S1.**
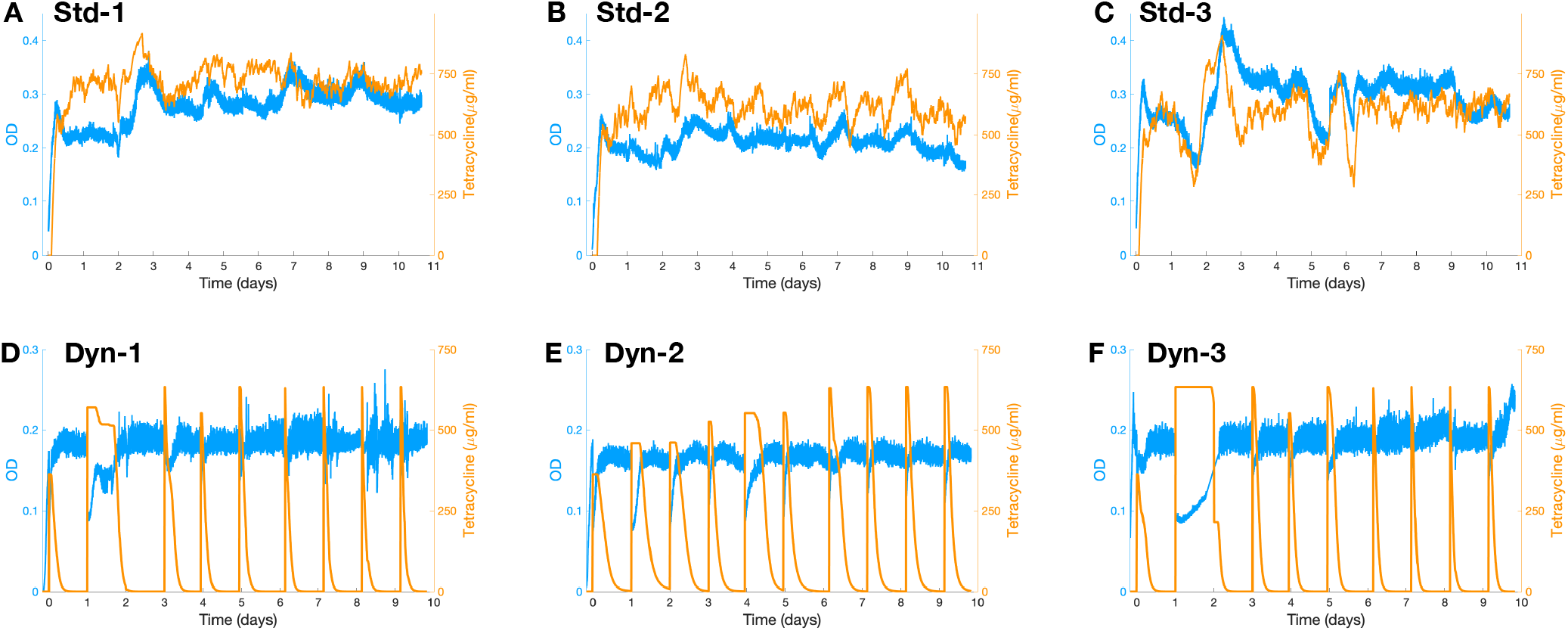
OD and tetracycline concentration over the course of the experiment. Optical densities and tetracycline concentrations measured over time for **A-C** Steady populations and **D-F** Dynamic populations. Populations Dyn-1 and Dyn-2 recovered slowly from the second exposure and skipped the third exposure.

**Figure S2.**
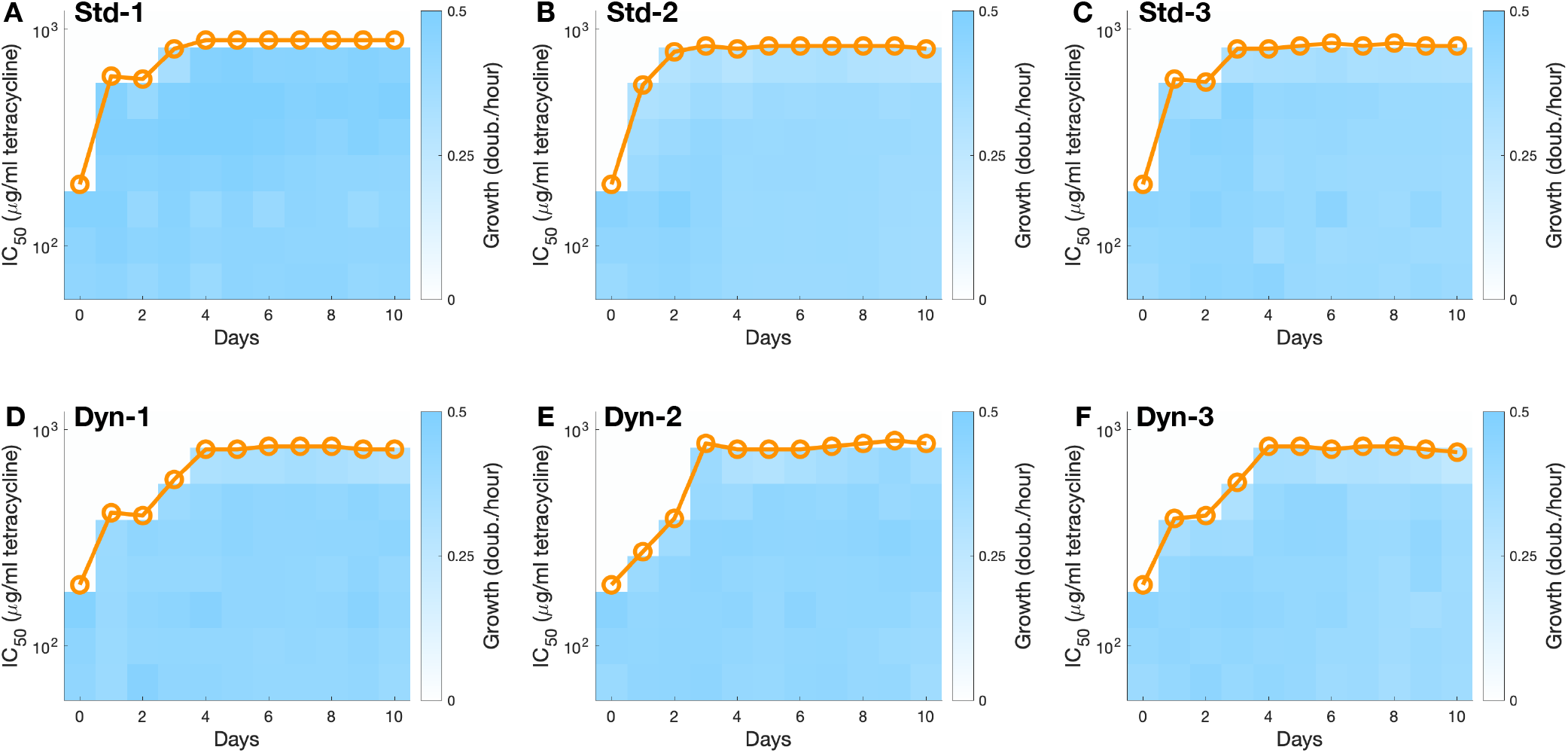
Progression of tetracycline resistance in each population. **A-F**. Samples from each population in each day were grown in tetracycline concentrations picked from a gradient, and growth rates were determined in each case. Circles indicate the drug concentration at which growth is reduced to half (IC_50_ concentration).

**Figure S3.**
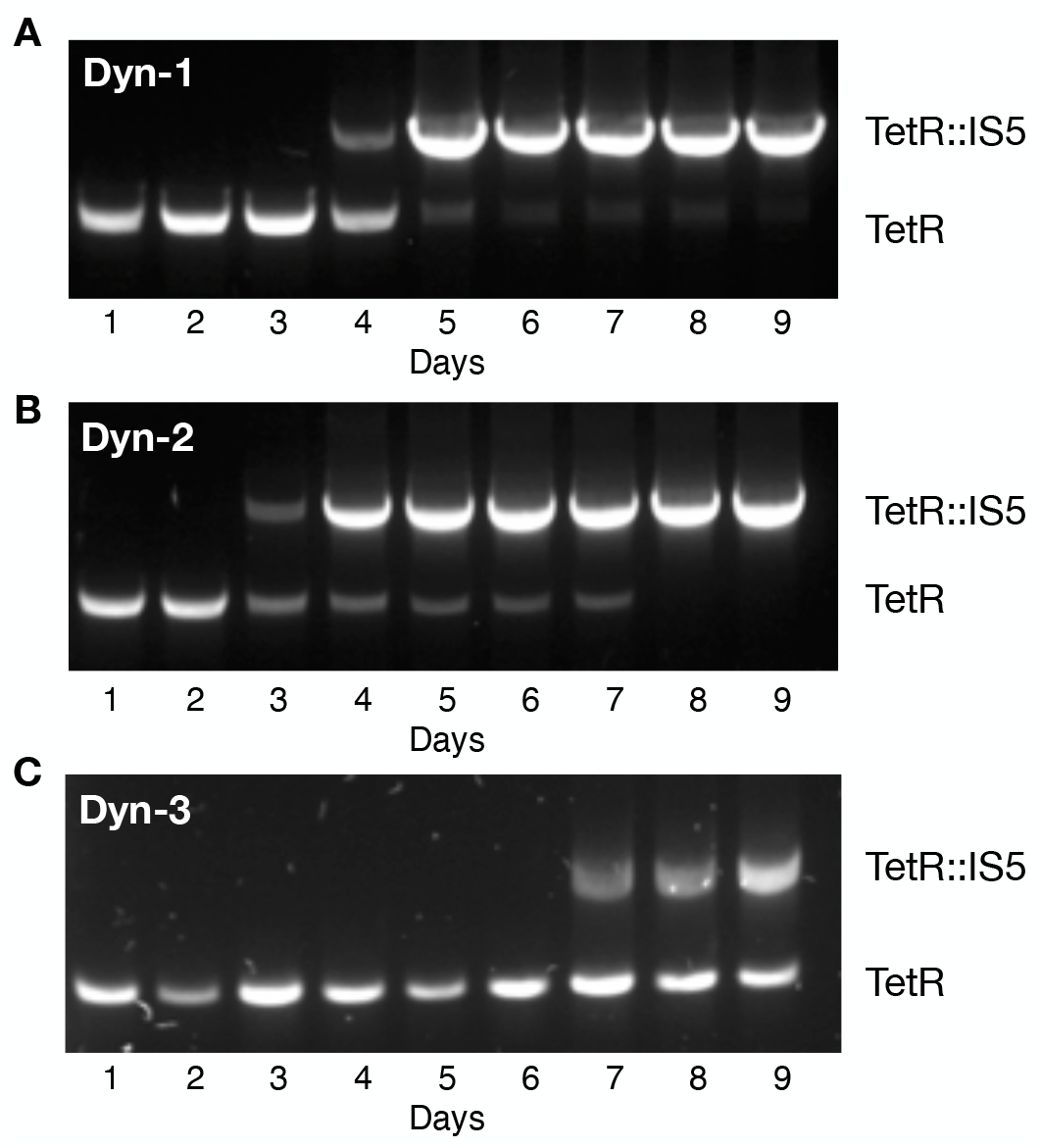
Prevalence of *IS5* insertions in the *tetR* gene over the course of the experiment. PCR amplification of the *tet* operon showing acquisition of transposon insertion in each Dynamic population. We quantified the relative intensity of the bands to estimate the prevalence of TetR transposon insertions in each population in each day.

**Figure S4.**
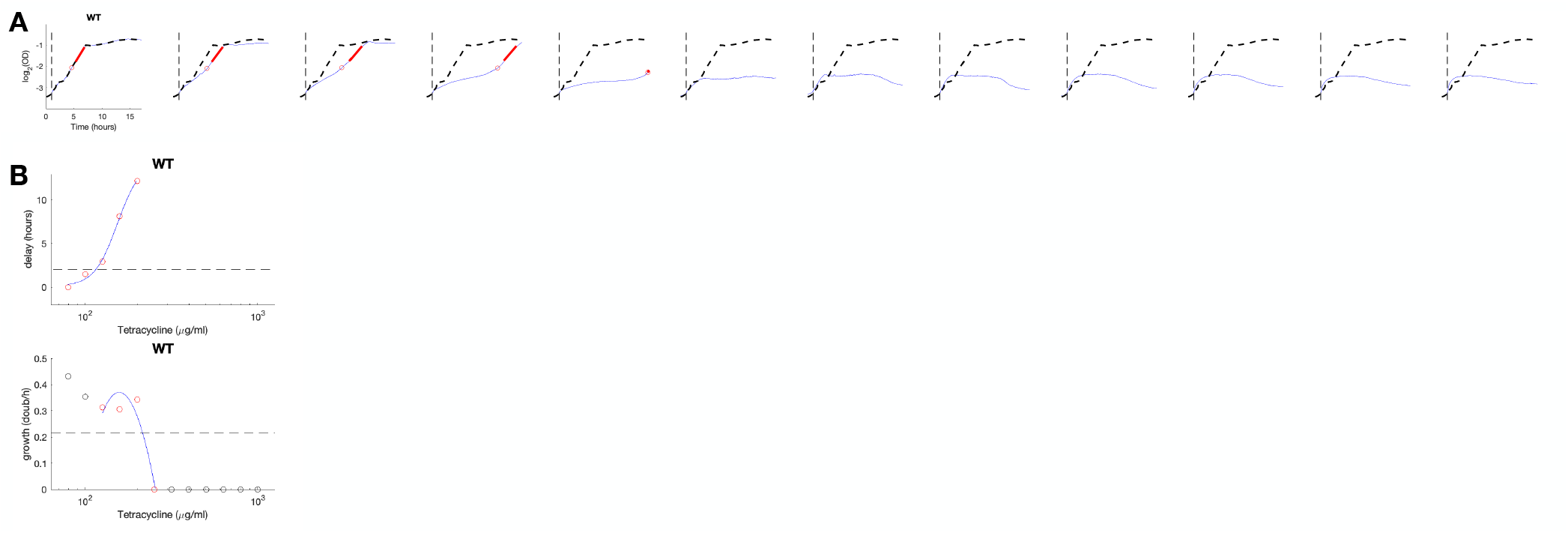
Calculation of steady-state and dynamical resistances for the WT ancestral strain. **A**. Growth curves of the WT ancestral strain. Plots are log2(OD) over 20 hours. Vertical dashed line indicates addition of tetracycline picked from a gradient of zero in the first column then 100 to 1000 μg/ml in columns 2 to 12. The corresponding growth curve with zero drug is included in a dashed line for comparison. The point at which the culture reaches a doubling is indicated, as well as the linear fit during steady-state growth following recovery. **B**. Dynamic resistance is calculated as the drug concentration that causes a 2-hour delay in the time to reach one doubling following drug exposure. Steady-state resistance is calculated as the drug concentration that reduces steady-state growth to half of the growth rate under no drug. Curves were fit with second order polynomials.

**Figure S5.**
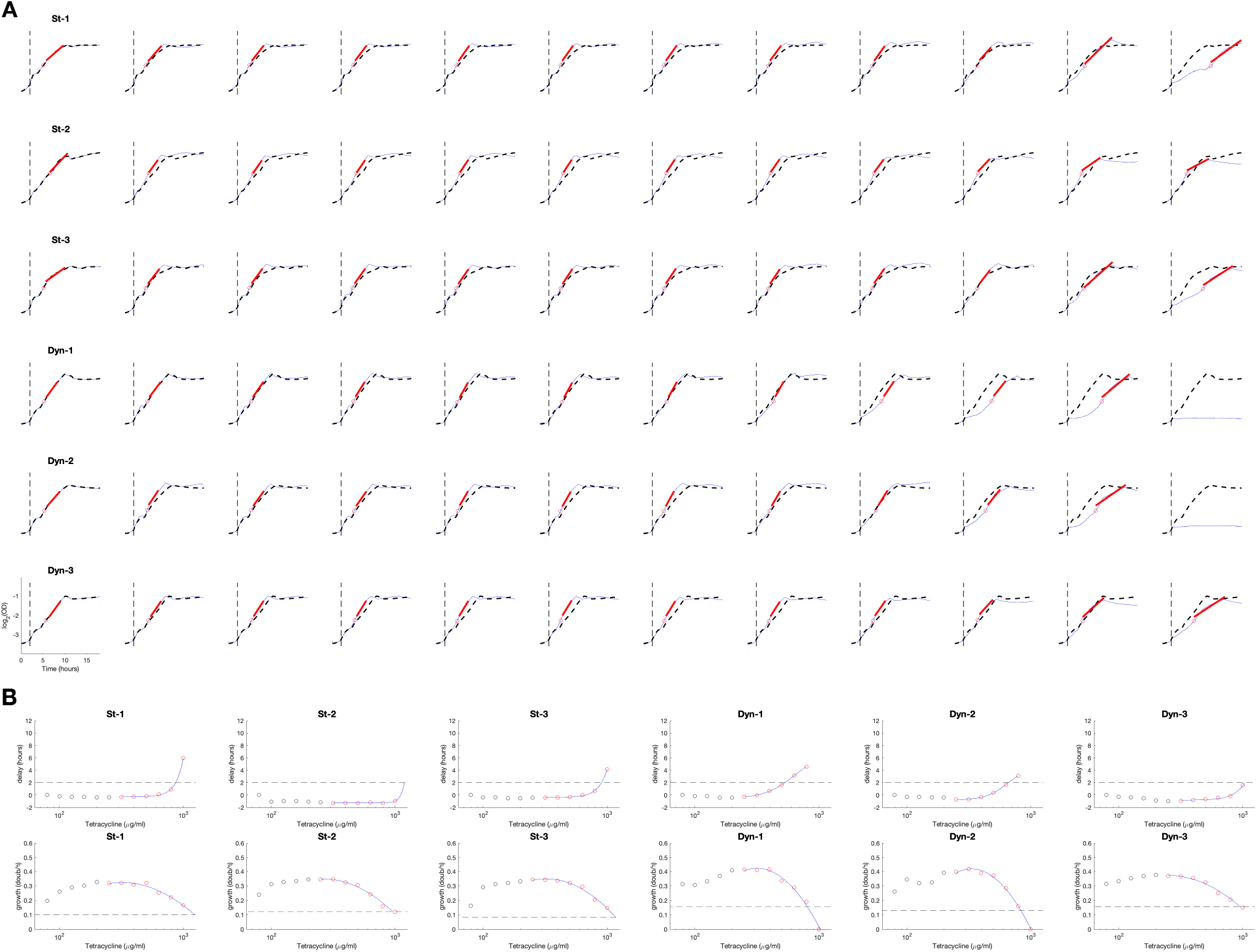
Calculation of steady-state and dynamical resistances for final evolved populations. **A**. Growth curves of the final populations. Plots are log2(OD) over 20 hours. Vertical dashed line indicates addition of tetracycline picked from a gradient of zero in the first column then 100 to 1000 μg/ml in columns 2 to 12. The corresponding growth curve with zero drug is included in a dashed line for comparison. The point at which the culture reaches a doubling is indicated, as well as the linear fit during steady-state growth following recovery. **B**. Dynamic resistance is calculated as the drug concentration that causes a 2-hour delay in the time to reach one doubling following drug exposure. Steady-state resistance is calculated as the drug concentration that reduces steady-state growth to half of the growth rate under no drug. Curves were fit with second order polynomials.

**Figure S6.**
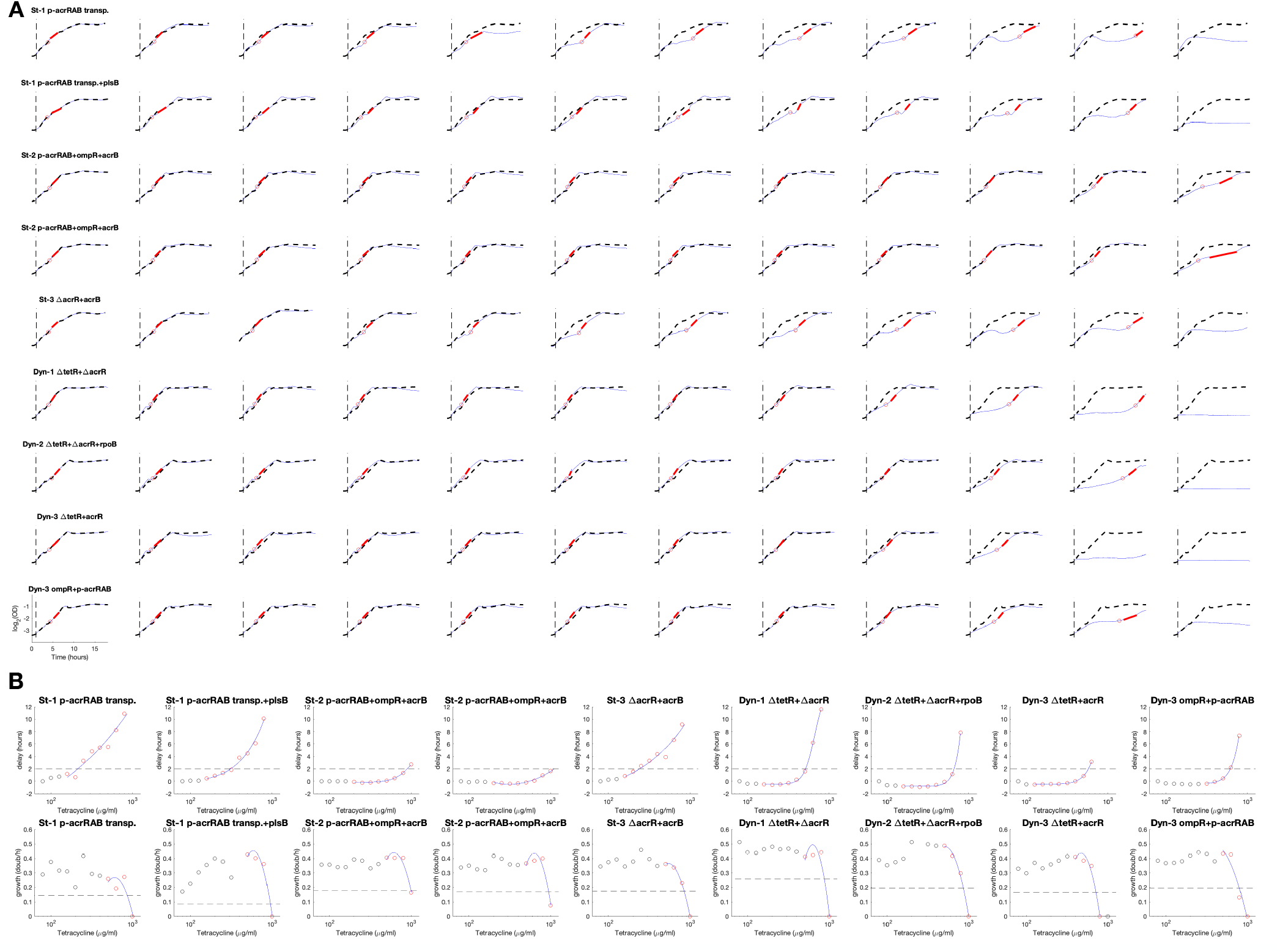
Calculation of steady-state and dynamical resistances for select mutants. **A**. Growth curves of select isolates picked from the final populations. Plots are log2(OD) over 20 hours. Vertical dashed line indicates addition of tetracycline picked from a gradient of zero in the first column then 100 to 1000 μg/ml in columns 2 to 12. The corresponding growth curve with zero drug is included in a dashed line for comparison. The point at which the culture reaches a doubling is indicated, as well as the linear fit during steady-state growth following recovery. **B**. Dynamic resistance is calculated as the drug concentration that causes a 2-hour delay in the time to reach one doubling following drug exposure. Steady-state resistance is calculated as the drug concentration that reduces steady-state growth to half of the growth rate under no drug. Curves were fit with second order polynomials.

**Figure S7.**
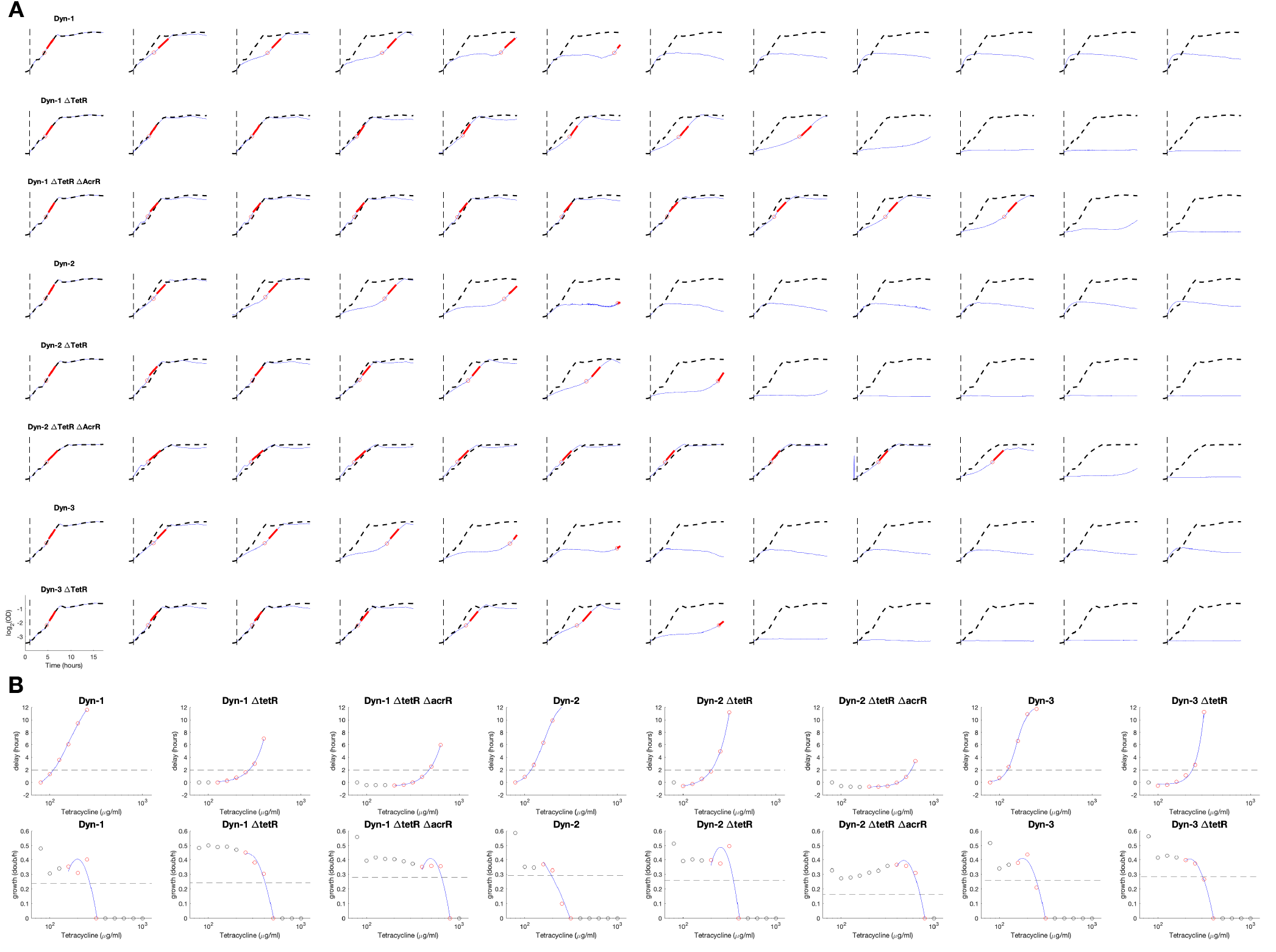
Calculation of gains in steady-state and dynamical resistances upon loss of TetR function. **A**. Growth curves of isolates picked in the same day, with and without transposon insertion in TetR, for each Dynamic population. Further acquisitions of AcrR mutations are also included. Plots are log2(OD) over 20 hours. Vertical dashed line indicates addition of tetracycline picked from a gradient of zero in the first column then 100 to 1000 μg/ml in columns 2 to 12. The corresponding growth curve with zero drug is included in a dashed line for comparison. The point at which the culture reaches a doubling is indicated, as well as the linear fit during steady-state growth following recovery. **B**. Dynamic resistance is calculated as the drug concentration that causes a 2-hour delay in the time to reach one doubling following drug exposure. Steady-state resistance is calculated as the drug concentration that reduces steady-state growth to half of the growth rate under no drug. Curves were fit with second order polynomials.

